# B-PPI: A Cross-Attention Model for Large-Scale Bacterial Protein-Protein Interaction Prediction

**DOI:** 10.64898/2025.12.23.696145

**Authors:** Chen Agassy, Bruria Samuel, Shahar Mayo, Asaf Zorea, David Burstein

## Abstract

Protein-protein interactions (PPIs) are essential for the study of cellular function, yet computational prediction of bacterial PPIs remains limited. Most existing methods are trained on human data, reducing their applicability to bacterial systems. Here, we present B-PPI, a computational tool specifically designed for bacterial PPI prediction. B-PPI leverages embeddings from ProstT5, a structure-aware protein language model, and a cross-attention mechanism to capture residue-level inter-protein relationships. To facilitate training, we constructed B-PPI-DB, a large-scale bacterial PPI dataset derived from STRING, comprising 202,829 positive and negative interactions across 2,646 taxa with a 1:10 positive-to-negative ratio. We benchmarked B-PPI against TT3D, a state-of-the-art model trained on human PPI, which was previously evaluated on bacterial PPIs. B-PPI achieved substantially higher performance on bacterial data (AUPRC 0.926±0.006 vs. 0.230±0.005 and F1 0.866±0.007 vs. 0.299±0.005) with faster runtime. We further demonstrate that the model adapts to unseen bacterial interactions with minimal fine-tuning. Together, B-PPI and B-PPI-DB address a critical gap in computational microbiology, offering a framework for bacterial PPI prediction and a data resource for benchmarking and developing new tools in the field.

## Introduction

Protein-protein interactions (PPIs) are fundamental to cellular physiology, governing processes ranging from signal transduction to metabolic regulation. In bacteria, resolving these interaction networks is particularly valuable for assigning functions to hypothetical proteins, reconstructing complex metabolic pathways, and understanding mechanisms of virulence and pathogenicity [1–4]. Since a large proportion of bacterial genomes remains functionally uncharacterized, computational PPI prediction can serve as a crucial first step in decoding these molecular systems.

Despite their biological importance, computational tools for rapid bacterial PPI prediction remain limited. While structure-based methods such as AlphaFold-Multimer [5] and the recently released AlphaFold3 [6] can model protein complexes with high accuracy, their computational intensity makes them impractical for large-scale screening. Consequently, there is a clear need for rapid, scalable frameworks that can filter and prioritize high-confidence interaction candidates in bacteria for subsequent structural validation.

Most existing rapid prediction tools, including D-SCRIPT [7], PIPR [8], XCapT5 [9] and TT3D [10], were developed primarily for eukaryotes. These models are typically trained on human data, which limits their generalizability to bacterial systems, likely due to the vast evolutionary distance and sequence divergence between prokaryotes and eukaryotes. When applied to bacteria, human-trained models show reduced performance [10], highlighting the necessity for tools optimized specifically for bacterial proteomes.

A further challenge in the field is the prevalence of artificially balanced training datasets (1:1 positive-to-negative ratio). While balancing data simplifies the learning process, it fails to reflect the reality of biological environments, where true interactions are sparse. Models trained on such distributions often yield inflated performance metrics that do not translate to real-world screening scenarios. To address this, recent studies have moved toward a 1:10 ratio for training and evaluation, a setting that provides a more rigorous test of a model’s ability to distinguish rare positive interactions from a background of negatives [10].

Many of those rapid prediction tools use protein Language Models (pLMs) embeddings as input to their models. Such pLMs have emerged as a powerful paradigm for protein representation, learning from vast sequence databases [11]. However, standard pLMs typically rely solely on amino acid sequences, potentially missing explicit structural signals that govern binding. ProstT5 [12] overcomes this limitation by adopting a “bilingual” training approach, learning from both amino acid sequences and Foldseek-derived 3Di representation, a sequence format that discretizes 3D structural features [13]. This allows the generation of embeddings that are inherently structure-aware, bridging the gap between rapid sequence-based processing and the structural precision required for interaction prediction.

To leverage these advances for bacterial systems, we curated B-PPI-DB, a comprehensive bacterial PPI dataset with a 1:10 positive-to-negative ratio. The positive interactions derived strictly from high-confidence, experimentally supported interactions in the STRING database [14]. By filtering out lower-confidence predictions and removing similar interactions, B-PPI-DB provides a robust benchmark tailored to bacterial systems.

Building on this resource, we introduce B-PPI, a screening tool specifically designed for bacterial PPI prediction. B-PPI utilizes ProstT5 embeddings to input implicitly encoded structural context into a bidirectional cross-attention transformer architecture. This design allows the model to capture complex residue-level inter-protein relationships. Benchmarked under identical conditions, B-PPI demonstrates superior performance over TT3D, a state-of-the-art human-centric model applied to bacteria, establishing a scalable framework for bacterial PPI prediction that can be adapted using minimal fine-tuning to bacterial species or interactions unseen during training.

## 2. Methods

### 2.1 Dataset Construction

Positive bacterial PPIs were obtained from the STRING database (version 12) [14]. To ensure high reliability, we selected interactions exclusively based on experimental evidence. Specifically, we retained interactions meeting one of two strict criteria: (1) a score ≥ 900 in the experiments channel; or (2) a score ≥ 900 in the experiments transferred channel, provided they were corroborated by a score ≥ 900 in the databases channel. This secondary criterion assumes that interactions conserved across taxa and annotated in curated bacterial databases represent homologous binding relationships in bacteria.

For out-of-distribution evaluation, we utilized the *H. pylori* benchmark dataset [15], consistent with the evaluation protocol of XCapT5 [9]. This set consists of 2,878 positive and negative interactions among *H. pylori* proteins. The positive interactions were validated via yeast two-hybrid assays [16]. To strictly prevent data leakage, all *H. pylori* proteins and their corresponding interactions were excluded from B-PPI-DB.

To remove redundancy and prevent homology-driven leakage, all bacterial proteins were clustered using MMseqs2 [17] at a 40% sequence identity threshold (--min-seq-id 0.4 -c 0.8 --cov-mode 0), following the protocol of D-SCRIPT [7]. Two interactions (*A, B*) and (*C, D*) were defined as redundant if protein *A* clustered with *C* and protein *B* clustered with *D* (or vice versa). We removed all redundant interactions from both positive and negative sets, retaining only the pair with the highest combined STRING score for each cluster combination in the positive set.

Negative examples were generated by randomly pairing proteins from the unique bacterial protein pool, ensuring that no sampled pair appeared in STRING as interacting at any confidence level. We applied a 1:10 positive-to-negative ratio, consistent with recent benchmarks such as TT3D [10]. While this ratio is less extreme than real biological interactomes, it provides a challenging yet computationally tractable setup for training and evaluation.

The resulting dataset, B-PPI-DB, comprises 202,829 total pairs (18,439 positive and 184,390 negative) involving 19,810 unique proteins across 2,646 bacterial taxa. B-PPI-DB is released as a community resource to facilitate future benchmarking and tool development.

### 2.2 Protein Representations

To capture both sequence semantics and structural features, each protein was represented using embeddings from ProstT5 [12]. Embeddings were extracted in half-precision (FP16) using the model available on the Hugging Face Hub [18]. For each protein of length *L*, we extracted residue-level embeddings, resulting in variable-length feature matrices of dimension *L* × 1024, which served as the direct input to our model.

### 2.3 Model Architecture

B-PPI employs a bidirectional cross-attention mechanism designed to capture inter-protein interaction signals and estimate interaction probability between protein pairs based on their residue-level embeddings (see Figure 1).

**Figure 1:**
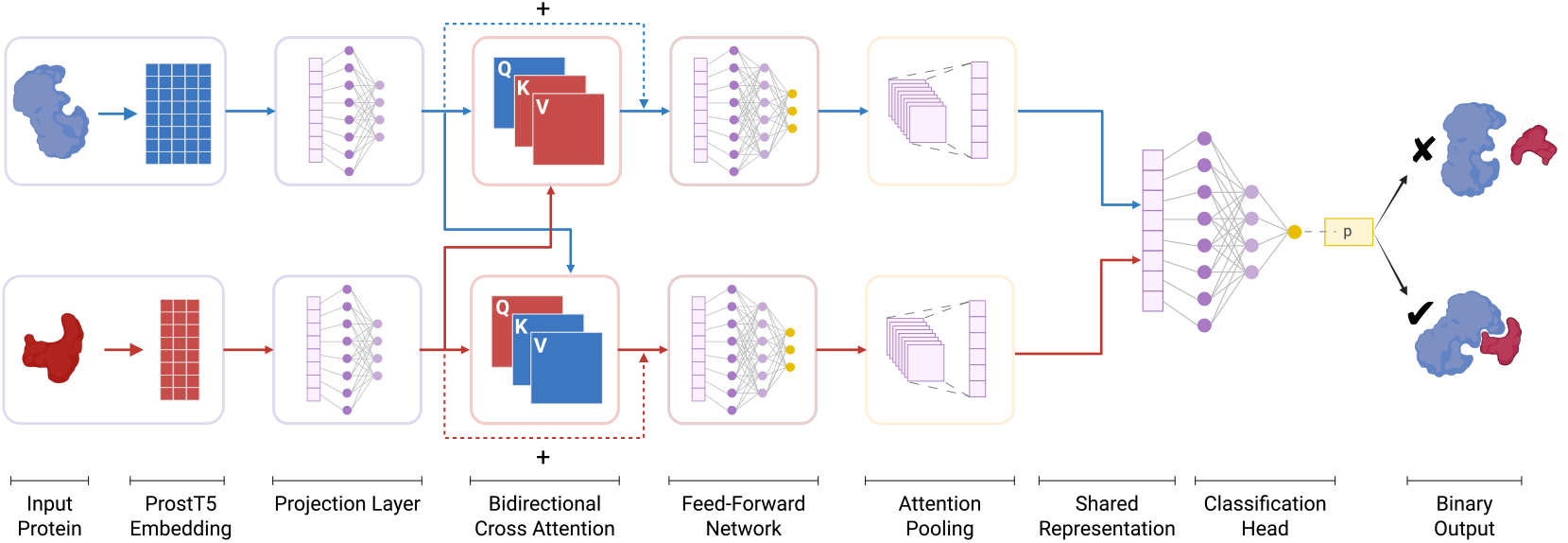
B-PPI Model Architecture. Input sequences are converted to ProstT5 embeddings, projected, and processed via bidirectional cross-attention with residual connections (Q: queries from self, K/V: keys/values from partner). The resulting features pass through a feed-forward network and attention pooling before being merged into a symmetric representation for the final classification layer, which outputs the predicted interaction probability, *p*.

**Figure 2.**
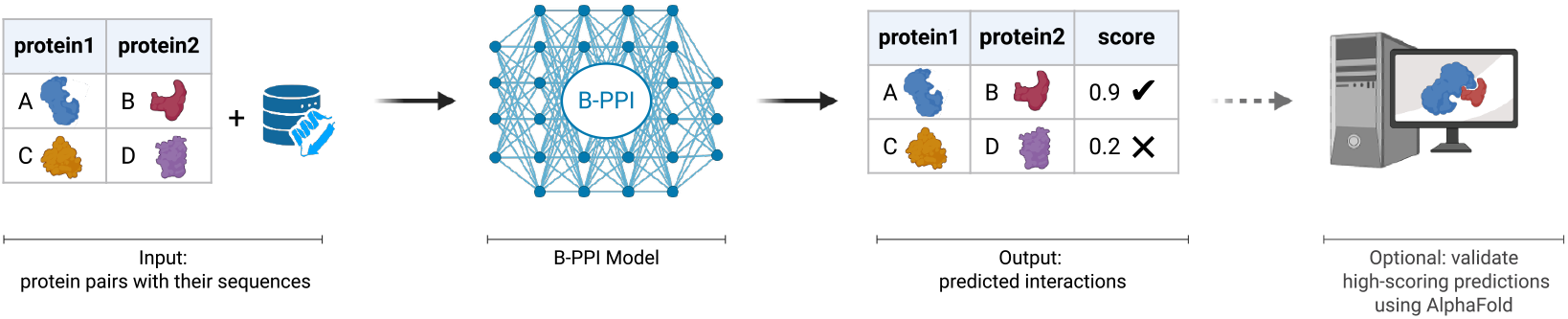
Illustration of the B-PPI workflow. Given a list of protein pairs and their corresponding sequences, the B-PPI model outputs the predicted interaction probabilities. High-confidence pairs can be validated with structural modeling tools such as AlphaFold3.

### 2.3.1 Input Projection

Let a protein pair be denoted as *P*_*A*_ and *P*_*B*_ with lengths *L*_*A*_ and *L*_*B*_, respectively. They are represented by embedding matrices 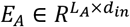 *and* 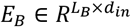 derived from ProstT5, where *d*_*in*_ = 1,024. To map these high-dimensional features to a compact latent interaction space of dimension *d*, we apply a learnable linear projection 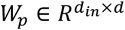:

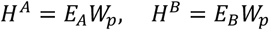

#### 2.3.2 Bidirectional Cross-Attention

To capture the dependencies between the two proteins, we employ a bidirectional Multi-Head Cross-Attention (MHCA) mechanism with *h* = 2 heads.

For the update of protein *P*_*A*_, the model treats *H*^*A*^ as the query source and *H*^*B*^ as the context (key/value) source. Normalized inputs are used to generate the attention components:

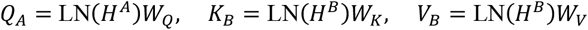

where LN denotes Layer Normalization. The attention weights are computed using the scaled dot-product attention:

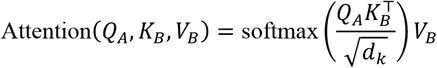

where *d*_*k*_ is the dimension per head. By adding the attention output to the original embedding *H*^*A*^ via a residual connection, the model fuses inter-protein interaction signals with the intra-protein context:

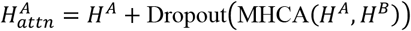

This process is performed symmetrically for protein *P*_*B*_ (where *P*_*B*_ queries *P*_*A*_). This block utilizes a dropout rate of 0.1 to prevent overfitting.

#### 2.3.3 Gated Feed-Forward Network

Following the attention mechanism, the representations are refined using a Feed-Forward Network (FFN) augmented with Gated Linear Units (GLU). This mechanism allows the model to selectively filter information by learning a multiplicative gate.

The FFN first expands the embedding dimension *d* to an intermediate width of 4,096 via a linear projection. Let *H*_*in*_ be the input. The projection yields a vector *V* ∈ *R*^4096^:

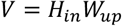

This vector *V* is split into two halves, *A* and *B*, each of dimension 2,048. The Gated Linear Unit operation is then applied, where *A* serves as the content and *B* serves as the gate:

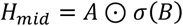

Finally, the filtered representation *H*_*mid*_ is projected back to the original dimension *d* and added to the input via a residual connection:

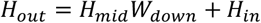

This block utilizes a dropout rate of 0.25 after both the up-projection and the down-projection to prevent overfitting.

#### 2.3.4 Attention Pooling

To transition from residue-level features to a pair-level interaction score, the variable-length residue embeddings *H* ∈ *R*^*L*×*d*^ are aggregated into fixed-size vectors using an Attention Pooling layer. For each protein, a learnable weight vector *w*_*a*_ and a projection matrix *W*_*a*_ are used to calculate a scalar importance score *α*_*i*_ for every residue *i*:

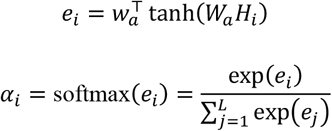

The final global representation *v* ∈ *R*^*d*^ is the weighted sum of the residue embeddings:

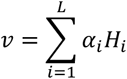

#### 2.3.5 Shared Representation

Since protein-protein interactions are physically undirected (i.e., the interaction *A* ↔ *B* is equivalent to *B* ↔ *A*), the model must be invariant to input order. We enforce this symmetry by constructing a joint representation *z* that concatenates the element-wise sum and absolute difference of the pooled vectors *v*_*A*_ and *v*_*B*_:

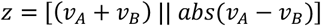

where || denotes vector concatenation.

#### 2.3.6 Classification Head

Finally, *z* is passed through a Multi-Layer Perceptron (MLP) to compute the interaction probability. The MLP consists of two hidden layers (sizes 256 and 64) with ReLU activation and dropout (*p* = 0.25), followed by a linear output layer and a Sigmoid activation:

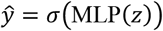

where ŷ ∈ [0,1] represents the predicted probability of interaction.

### 2.4 Training and Optimization Strategy

To address the class imbalance inherent in PPI datasets, where non-interacting pairs vastly outnumber interacting ones, we trained B-PPI using Focal Loss [19] rather than standard cross-entropy. Focal Loss dynamically scales the loss based on the confidence of the prediction, focusing the model on hard-to-classify examples. The loss function is defined as:

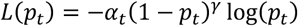

where *p*_*t*_ is the model’s estimated probability for the true class. We set the balancing factor α to 0.9 to significantly upweight positive examples and the focusing parameter γ to 2 to reduce the penalty for easily classified negatives. This combination ensures that the gradient is driven by difficult, ambiguous examples rather than the overwhelming number of easy negatives.

Optimization was performed using the AdamW algorithm [20] with a learning rate of 5 × 10^−5^ and weight decay of 1 × 10^−4^. Training proceeded in batches of 64 pairs for 10 epochs. Hyperparameters were selected via random search to maximize the Area Under the Precision-Recall Curve (AUPRC) on the validation set.

### 2.5 Cross-Validation

To test the ability of B-PPI to predict STRING experimental interactions, we employed five-fold cross-validation. We selected five folds as a balance between computational efficiency and reliable estimation of model performance. In each fold, an independent 20% of the data was reserved for testing, while the remaining 80% was split into 99% for training and 1% for validation to facilitate early stopping.

To prevent information leakage and encourage the model to learn generalizable interaction principles, the cross-validation splits were cluster-aware at the interaction level, as described in Section 2.1. Specifically, interactions were considered redundant if both proteins belonged to clusters already represented in another pair. By removing redundant interactions across folds, the model is discouraged from memorizing specific interactions, ensuring more robust evaluation of its predictive performance.

### 2.6 Fine-Tuning

To assess the model’s adaptability to out-of-distribution data, we performed minimal fine-tuning using the *H. pylori* dataset [15]. We implemented a fine-tuning only to the classification head: the parameters for the ProstT5 projection, cross-attention mechanism, and FFN were frozen, and only the final MLP prediction head was updated.

Fine-tuning was conducted for 100 epochs using the AdamW optimizer with a learning rate of 1 × 10^<0^ and Binary Cross-Entropy (BCE) loss. The dataset was split into 50% for training the prediction head, 25% for validation and 25% for testing. This setup simulates a “low-resource” biological scenario, where a general model must be adapted to a specific bacterial species or interactions using only a limited set of experimentally validated interactions.

## 3. Results

### 3.1 B-PPI-DB

We constructed B-PPI-DB, a large-scale dataset comprising 202,829 non-redundant bacterial protein-protein interactions with a 1:10 positive-to-negative ratio (18,439 positive and 184,390 negative) across 2,646 taxa derived from STRING [14]. B-PPI-DB provides a challenging benchmark that better reflects the sparsity of biological interactomes compared to balanced datasets, and is released as a community resource to facilitate future benchmarking and tool development.

### 3.2 B-PPI Pipeline

We developed B-PPI to address the scarcity of computational tools specifically optimized for large-scale screening of bacterial protein-protein interactions. The model was trained and rigorously evaluated using B-PPI-DB.

To facilitate large-scale PPI screening, the B-PPI pipeline was designed as a streamlined workflow accepting FASTA sequences and a file specifying the candidate pairs to be tested. The model computes interaction probabilities and binary classifications for each pair, enabling researchers to prioritize high-confidence candidates for structural validation with tools like AlphaFold3 [6].

Central to this workflow is the B-PPI model, where input sequences are initially encoded using ProstT5

[12] to capture sequence and structural semantics and projected into a lower-dimensional latent space. These representations are subsequently processed by a bidirectional cross-attention mechanism with residual connections, which models residue-level dependencies by allowing each protein to dynamically attend to its partner. The resulting interaction signals are refined via a feed-forward network and aggregated via attention pooling, after which the pooled embeddings are combined into a symmetric joint representation to yield the predicted interaction probability.

### 3.3 Comparative Performance on B-PPI-DB

To rigorously evaluate B-PPI, we benchmarked it against TT3D [10], a state-of-the-art model originally trained on human interactions. TT3D employs a dual-input architecture, combining standard amino acid sequences with 3Di structural tokens [13] to explicitly integrate both sequence and structural information. In contrast, B-PPI leverages direct ProstT5 [12] embeddings, which implicitly encode both sequence semantics and latent structural context within a single, unified representation.

To ensure a strict sequence-based comparison that does not rely on experimental structures, the 3Di sequences required by TT3D were generated using ProstT5. This ensures that both models operate on identical information sources - derived solely from protein sequences - while utilizing different representation strategies.

Beyond predictive accuracy, B-PPI offers distinct computational advantages for large-scale screening. TT3D relies on sequential decoding to generate 3Di tokens, resulting in inference latencies of 0.6–2.5 seconds per protein. In contrast, B-PPI utilizes direct half-precision embedding extraction, which reduces computation time to approximately 0.1 seconds per protein on an NVIDIA RTX A6000 GPU [18]. This efficiency removes a critical bottleneck in the inference pipeline, positioning B-PPI as a scalable solution for large-scale PPI prediction.

We evaluated both models using the strict five-fold cross-validation framework described in the Methods section. TT3D predictions were computed on the exact same test partitions to ensure direct comparability. Table 1 summarizes the mean performance across folds, with metrics reported at the decision threshold that maximizes the F1 score. Detailed performance metrics for each individual fold are provided in Supplementary Tables S1 and S2.

**Table 1.** Performance metrics for B-PPI and TT3D (five-fold cross-validation). Metrics are reported as mean±standard deviation at the threshold maximizing the F1 score.

| Model | AUROC | AUPRC | Accuracy | Precision | Recall | F1 | Max F1 Threshold |
| --- | --- | --- | --- | --- | --- | --- | --- |
| **B-PPI** | **0.992**<br>$\pm 0.001$ | **0.926**<br>$\pm 0.006$ | **0.974**<br>$\pm 0.002$ | **0.823**<br>$\pm 0.013$ | **0.914**<br>$\pm 0.006$ | **0.866**<br>$\pm 0.007$ | **0.627**<br>$\pm 0.029$ |
| **TT3D** | 0.688<br>$\pm 0.005$ | 0.230<br>$\pm 0.005$ | 0.848<br>$\pm 0.003$ | 0.258<br>$\pm 0.004$ | 0.358<br>$\pm 0.012$ | 0.299<br>$\pm 0.005$ | 0.141<br>$\pm 0.014$ |

B-PPI consistently outperforms the TT3D across all evaluation criteria. Most notably, B-PPI achieved an AUROC improvement of over 0.3 and a nearly four-fold increase in AUPRC compared to TT3D. The ROC and Precision-Recall curves (Figure 3) further illustrate this performance gap. B-PPI maintains a high true positive rate across the entire false positive rate spectrum and retains high precision even at high recall levels. In contrast, TT3D’s precision degrades rapidly, reflecting the difficulty of transferring human-centric interaction rules to bacterial data.

**Figure 3.**
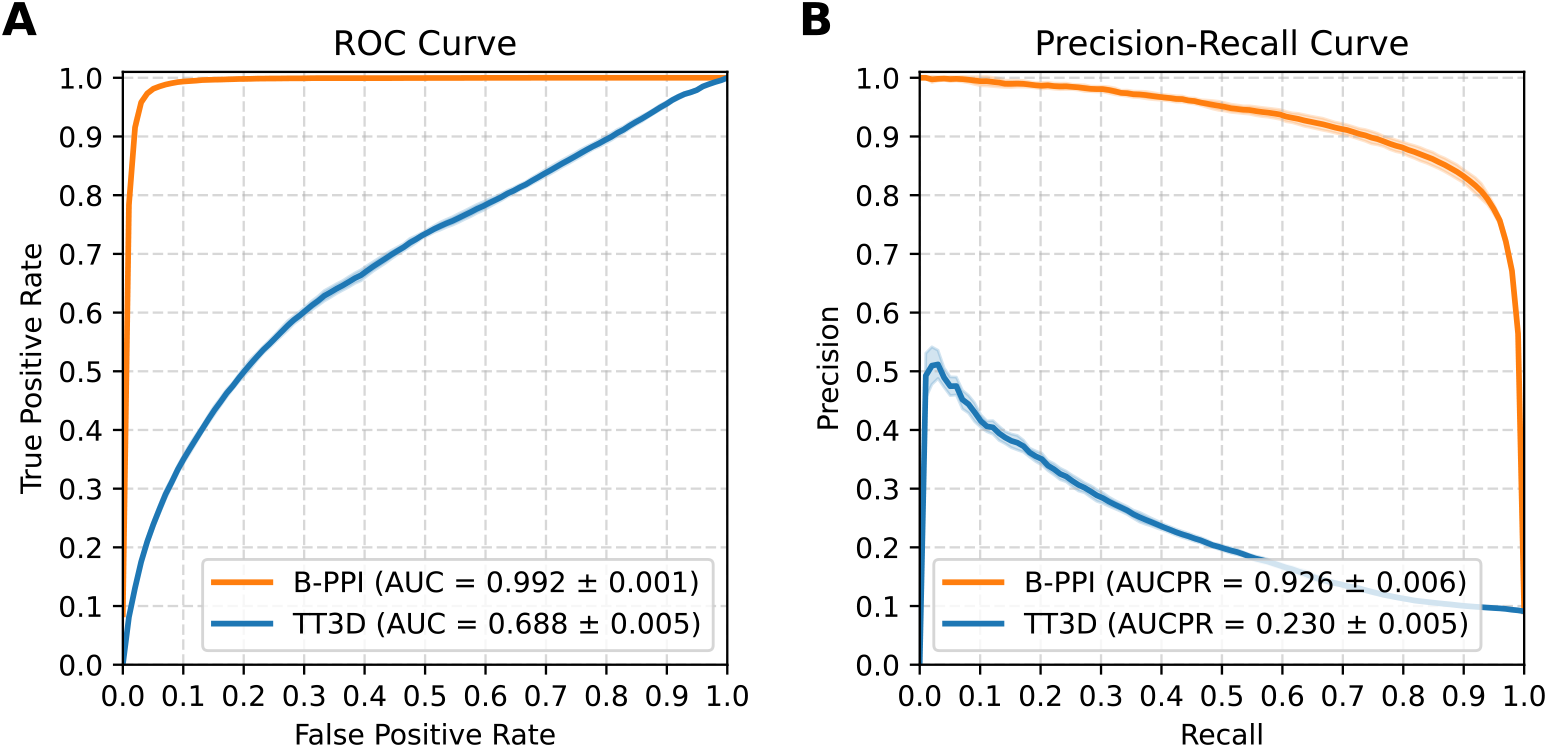
Performance comparison between B-PPI and TT3D across folds. **(A)** ROC curve comparison. **(B)** Precision-Recall curve comparison. B-PPI exhibits superior global discrimination and maintains precision at higher recall thresholds.

Furthermore, an analysis of the predicted probability distributions (Figure 4) reveals that TT3D assigns low probabilities to many true positive bacterial interactions, resulting in a substantial overlap with negative pairs. In contrast, B-PPI produces a clear separation between classes, which underpins its ability to minimize false positives in large-scale screening contexts.

**Figure 4.**
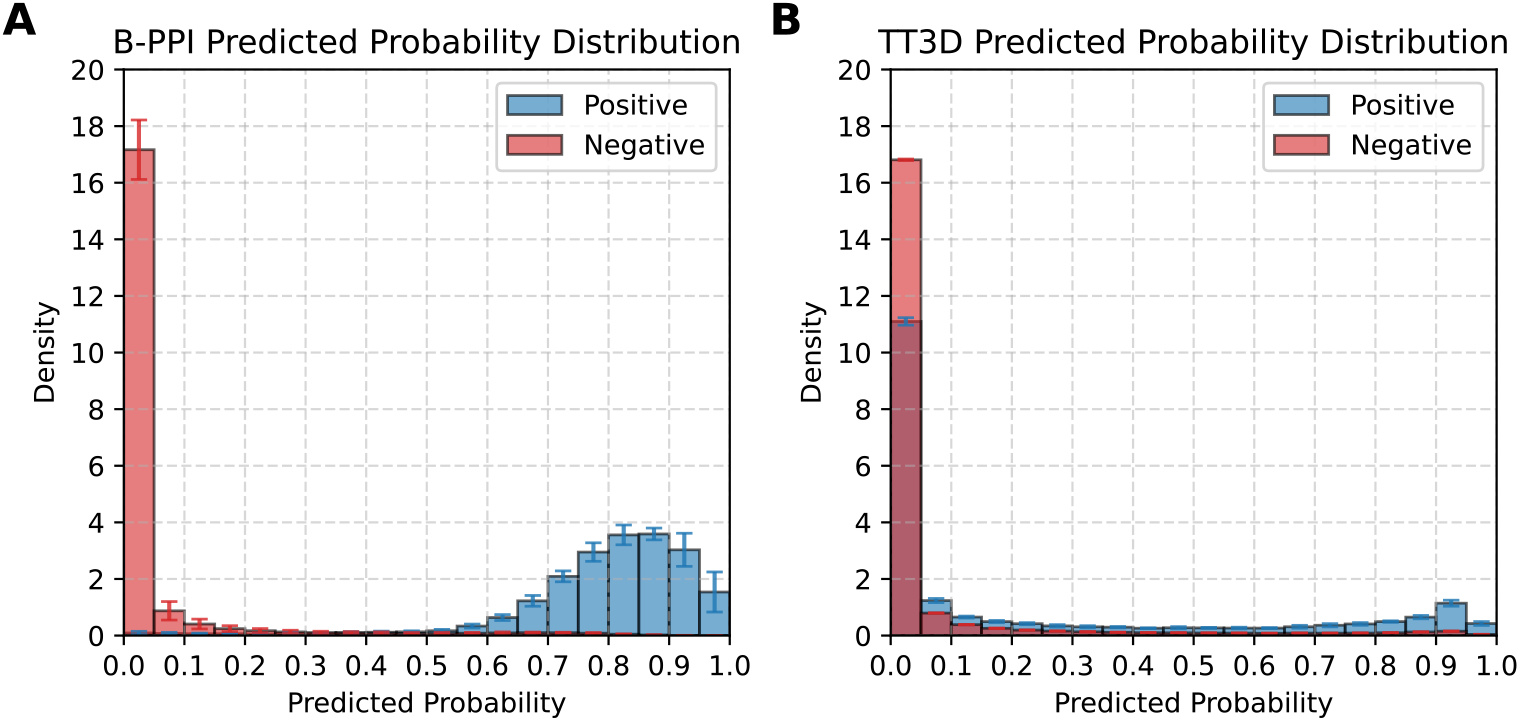
Predicted probability distributions across folds. **(A)** B-PPI predicted probability distribution (**B)** TT3D predicted probability distribution. B-PPI exhibits a clean separation between interacting (positive, in blue) and non-interacting (negative, in red) populations compared to TT3D.

This distinction is further corroborated by confusion matrix analysis provided in Supplementary Figure S3. At the decision threshold maximizing the F1 score, confusion matrices reveal that B-PPI achieves significantly higher counts of true positives and true negatives while simultaneously reducing false positives and false negatives relative to TT3D. This balanced reduction in misclassifications demonstrates that B-PPI’s superior performance is not driven by biasing towards the majority class, but by genuinely improved discrimination of interaction features.

### 3.4 Fine-Tuning on an External Dataset

Beyond the five-fold cross-validation on B-PPI-DB, we further evaluated whether the trained model could generalize to out-of-distribution interactions, specifically, proteins from taxa entirely unseen during training.

To this end, we excluded all *H. pylori* proteins from the B-PPI-DB during construction (see Methods) and utilized the *H. pylori* benchmark dataset [15] for external evaluation, following a similar evaluation procedure to that used in XCapT5 [9]. This dataset consists of 1,420 positive and 1,458 negative interactions.

For this specific evaluation, a base model was trained on 95% of the B-PPI dataset, with the remaining 5% reserved for validation (2.5%) and testing (2.5%). When applying this STRING-trained model to the *H. pylori* dataset in a zero-shot setting (without additional training), performance dropped notably to an AUPRC of 0.4 and an F1 score of 0.66. This decline reflects the fundamental challenge of out-of-distribution prediction, as the specific interaction patterns in this dataset are probably not sufficiently represented within the training data distribution.

To assess adaptability, we performed minimal fine-tuning (as described in Methods) using half of the *H. pylori* dataset (1,439 labeled examples). This adaptation resulted in a substantial performance improvement, achieving an AUPRC of 0.81 and an F1 score of 0.77.

These findings demonstrate that B-PPI can effectively serve as a foundation model for bacterial PPI prediction. While the model requires adaptation when applied to interactions that are distant from the training distribution, the lower layers of the network encode robust feature representations. This adaptability highlights the potential of B-PPI for practical applications where large-scale experimental data are typically unavailable.

## 4. Discussion

The development of B-PPI and the construction of B-PPI-DB represent a targeted effort to address the scarcity of computational tools optimized for rapid bacterial protein-protein interaction prediction. By combining a domain-specific training strategy with a bidirectional cross-attention architecture that leverages structure-aware ProstT5 [12] embeddings, this work aims to improve the scalability and accuracy of PPI screening in prokaryotic systems.

Our benchmarking results indicate that B-PPI outperforms TT3D [10], a state-of-the-art model trained on human data, across all evaluation metrics when tested on bacterial data. This performance gap suggests that interactions derived primarily from human data do not generalize well enough to bacterial proteomes, highlighting the utility of the B-PPI-DB dataset, which aggregates over 200,000 non-redundant positive and negative bacterial interactions.

Unlike methods often validated on artificially balanced datasets, B-PPI was trained and tested using a 1:10 positive-to-negative ratio. While this ratio is still less sparse than actual biological interactomes, it provides a more rigorous estimate of discriminative power than balanced benchmarks. The high AUPRC achieved under these conditions indicates that the model effectively minimizes false positives in this controlled setting, a critical requirement for screening applications.

Furthermore, the external evaluation on the *H. pylori* dataset [15] offers insight into the model’s adaptability. The observation that the pre-trained model’s performance dropped in a zero-shot setting but recovered significantly after fine-tuning only the final classification layers, suggests that the lower layers of the network encode robust, transferable representations. This indicates that B-PPI has potential as a foundational framework that can be adapted to distinct interaction distributions with limited labeled data.

Despite these promising results, several limitations must be acknowledged. First, the B-PPI-DB dataset relies on STRING [14], which introduces specific biases toward well-studied model organisms and conserved pathways. In Addition, positive interactions in STRING often conflate direct physical contacts with co-complex membership. Second, while the 1:10 training ratio improves upon balanced baselines, it does not fully represent the extreme sparsity of proteomes, where non-interacting pairs vastly outnumber true interactions. Therefore, in real-world proteomic scans, the false discovery rate may be higher than estimated in this study. Finally, as evidenced by the initial drop in performance on the *H. pylori* data, the model generalizes less effectively to taxa or interaction patterns that diverge from the training distribution without explicit adaptation.

Given these factors, B-PPI is best utilized as a high-efficiency filtration tool, rather than a standalone confirmation method. With rapid inference, the model is well-suited to scan bacterial proteomes (or metaproteomes) and prioritize high-confidence candidates. These candidates can then be subjected to high-resolution structural validation using computationally intensive tools such as AlphaFold3 [6]. Used in this manner, B-PPI provides a scalable and modular framework for PPI prediction in bacteria.

Moreover, the B-PPI-DB dataset serves as a valuable community resource and a rigorous benchmark for future developments in the field.

In summary, B-PPI lays the foundation for future methods specifically tailored to bacterial networks, demonstrating the distinct value of combining domain-specific bacterial data with advanced protein language model embeddings and cross-attention architecture.

## Acknowledgement

We thank Ella Rannon for her valuable feedback during the research and reading of the manuscript, and Rafael Pinilla-Redondo for stimulating discussion leading to the conception of this paper.

## Code and Data Availability

The models and scripts developed in this study are available on GitHub: https://github.com/burstein-lab/B-PPI.git.

The B-PPI-DB benchmarking database is available via Zenodo: https://doi.org/10.5281/zenodo.17927352.

