## Supplemental Tables and Figures for "B-PPI: A Cross-Attention Model for Large-Scale Bacterial Protein-Protein Interaction Prediction"

### Supplementary

#### 5-fold cross-validation evaluation results

| | AUROC | AUPRC | Accuracy | Precision | Recall | F1 | Threshold Maximizing F1 |
|---|---|---|---|---|---|---|---|
| 1 | 0.991 | 0.917 | 0.972 | 0.801 | 0.915 | 0.854 | 0.662 |
| 2 | 0.992 | 0.920 | 0.975 | 0.833 | 0.904 | 0.867 | 0.658 |
| 3 | 0.993 | 0.929 | 0.974 | 0.819 | 0.916 | 0.865 | 0.606 |
| 4 | 0.992 | 0.932 | 0.975 | 0.825 | 0.921 | 0.870 | 0.588 |
| 5 | 0.993 | 0.931 | 0.976 | 0.837 | 0.915 | 0.874 | 0.620 |
| Mean±Std | 0.992±0.001 | 0.926±0.006 | 0.974±0.002 | 0.823±0.013 | 0.914±0.006 | 0.866±0.007 | 0.627±0.029 |

**Table S1. Performance metrics for B-PPI on the five-fold cross-validation**. Metrics are reported at the threshold corresponding to maximum F1.

| | AUROC | AUPRC | Accuracy | Precision | Recall | F1 | Threshold Maximizin g F1 |
|---|---|---|---|---|---|---|---|
| 1 | 0.684 | 0.228 | 0.852 | 0.261 | 0.343 | 0.296 | 0.151 |
| 2 | 0.690 | 0.233 | 0.843 | 0.252 | 0.366 | 0.298 | 0.127 |

| | | | | | | | |
|---|---|---|---|---|---|---|---|
| 3 | 0.682 | 0.226 | 0.851 | 0.260 | 0.347 | 0.297 | 0.162 |
| 4 | 0.689 | 0.227 | 0.846 | 0.253 | 0.356 | 0.296 | 0.135 |
| 5 | 0.697 | 0.238 | 0.847 | 0.262 | 0.377 | 0.309 | 0.129 |
| Mean±Std | 0.688±0.005 | 0.230±0.005 | 0.848±0.003 | 0.258±0.004 | 0.358±0.012 | 0.299±0.005 | 0.141±0.014 |

**Table S2. Performance metrics for TT3D on the five-fold cross-validation**. Metrics are reported at the threshold corresponding to maximum F1.

#### Confusion Matrix Analysis

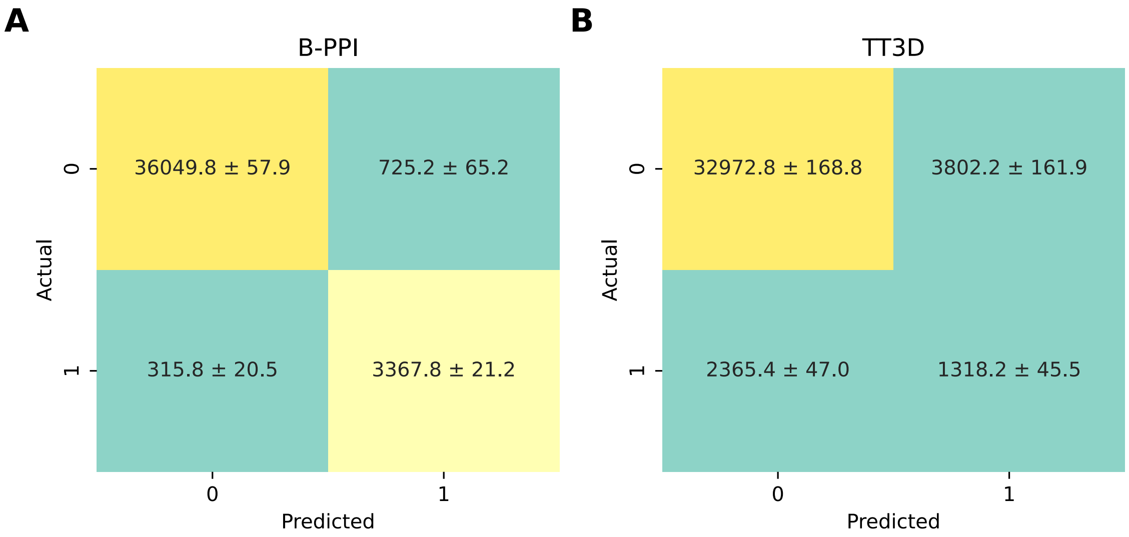


**Figure S3. Confusion matrices across folds at the decision threshold maximizing the F1 score.** (**A**) B-PPI confusion matrix. **(B)** TT3D confusion matrix. B-PPI correctly identifies more positives and negatives, reducing misclassifications compared to TT3D.
